# Model-based frequency-and-phase correction of ^1^H MRS data with 2D linear-combination modeling

**DOI:** 10.1101/2024.03.26.586804

**Authors:** Dunja Simicic, Helge J. Zöllner, Christopher W. Davies-Jenkins, Kathleen E. Hupfeld, Richard A. E. Edden, Georg Oeltzschner

**Affiliations:** Russell H. Morgan Department of Radiology and Radiological Science, The Johns Hopkins University School of Medicine, Baltimore, MD, United States; F. M. Kirby Research Center for Functional Brain Imaging, Kennedy Krieger Institute, Baltimore, MD, United States

**Keywords:** magnetic resonance spectroscopy, frequency-and-phase correction, spectral registration, two-dimensional linear-combination model

## Abstract

**Purpose:** Retrospective frequency-and-phase correction (FPC) methods attempt to remove frequency-and-phase variations between transients to improve the quality of the averaged MR spectrum. However, traditional FPC methods like spectral registration struggle at low SNR. Here, we propose a method that directly integrates FPC into a two-dimensional linear-combination model (2D-LCM) of individual transients (‘model-based FPC’). We investigated how model-based FPC performs compared to the traditional approach, i.e., spectral registration followed by 1D-LCM in estimating frequency-and-phase drifts and, consequentially, metabolite level estimates.

**Methods:** We created synthetic in-vivo-like 64-transient short-TE sLASER datasets with 100 noise realizations at 5 SNR levels and added randomly sampled frequency and phase variations. We then used this synthetic dataset to compare the performance of 2D-LCM with the traditional approach (spectral registration, averaging, then 1D-LCM). Outcome measures were the frequency/phase/amplitude errors, the standard deviation of those ground-truth errors, and amplitude Cramér Rao Lower Bounds (CRLBs). We further tested the proposed method on publicly available in-vivo short-TE PRESS data.

**Results:** 2D-LCM estimates (and accounts for) frequency-and-phase variations directly from uncorrected data with equivalent or better fidelity than the conventional approach. Furthermore, 2D-LCM metabolite amplitude estimates were at least as accurate, precise, and certain as the conventionally derived estimates. 2D-LCM estimation of frequency and phase correction and amplitudes performed substantially better at low-to-very-low SNR.

**Conclusion:** Model-based FPC with 2D linear-combination modeling is feasible and has great potential to improve metabolite level estimation for conventional and dynamic MRS data, especially for low-SNR conditions, e.g., long TEs or strong diffusion weighting.

## 1. Introduction

Proton magnetic resonance spectroscopy (^1^H MRS) is a non-invasive technique used to investigate metabolites in living tissue, particularly in the brain, with numerous applications in clinical practice and research^1,2^. The primary outcome measures of in-vivo ^1^H MRS experiments are concentration estimates for different metabolites in the measurement volume. A typical analysis workflow transforming raw data into quantitative concentration estimates requires three important phases: preprocessing, spectral modeling, and quantification^3^.

The data acquired during a conventional MRS experiment consist of multiple transients. It is common practice to combine them during preprocessing: since the signal is consistent between transients and the noise is random, coherent averaging generates a single spectrum with a sufficient signal-to-noise ratio (SNR) for subsequent modeling. In practice, experimental instabilities such as thermal B_0_ drift or participant motion cause frequency-and-phase variations between transients. If these effects are not corrected, the averaged spectrum may exhibit apparent linebroadening, resulting in greater modeling uncertainty due to reduced SNR and poorer spectral resolution^3–5^.

Retrospective frequency-and-phase correction (FPC) methods attempt to estimate and remove the frequency-and-phase variations between transients and therefore improve the quality of the averaged spectrum^3^. While active prospective frequency correction during the acquisition is possible^6,7^, it requires additional acquisitions or sequence modifications that are not available for all systems and does not address phase. In contrast, retrospective correction methods reduce frequency-and-phase variations post-hoc, i.e., during data analysis. Early implementations estimated the required corrections from selected singlet signals, for example, the residual water or total creatine peaks^10,13–15^. Improved methods use the full transient in the time (e.g., spectral registration^4^) or frequency domain (e.g., RATS^5^) and determine the necessary corrections by minimizing differences between the acquired transients. These methods generally outperform single-peak alignment in terms of robustness; but, at low SNR (e.g., SNR < 5 per transient), shot-to-shot alignment becomes challenging^5,16^. Furthermore, FPC’s performance is affected by residual water signals and lipid contamination, which create variable between-transient changes distinct from the frequency-and-phase drifts targeted by FPC^16^. Even though several methods attempt to minimize this effect by excluding the water/lipid region^17^ or filtering them prior to FPC^16^, correcting low-SNR data remains a challenge^5,16^.

Here, we propose a method that directly integrates FPC into a two-dimensional linear-combination model (2D-LCM) across individual transients (‘model-based frequency-and-phase correction’). 2D-LCM algorithms fit a set of spectra simultaneously, allowing the inclusion of relationships between these spectra in a single model. 2D models were initially proposed for 2D-resolved techniques (e.g., JPRESS)^18^ and have recently gained popularity due to an increased interest in advanced ^1^H MRS techniques such as relaxometry and diffusion-weighted^19^ or functional^20^ MRS, where experimental conditions are purposely modified between acquisitions to encode dynamic/temporal changes. This approach was shown to outperform the traditional averaging and 1D-LCM combined with the a posteriori fitting of a dynamic model, leading to improved precision and accuracy of estimated metabolite amplitudes^21^. Furthermore, we recently showed that 2D-LCM of a simplified (no frequency-and-phase variations) conventional synthetic MRS experiment offers comparable accuracy and precision to 1D-LCM of the averaged spectrum, with small benefits in uncertainty estimation^22^.

In this study, we explored whether 2D model-based correction of conventional MRS experiments offers comparable or superior (at low SNRs) performance to the standard approach (spectral registration followed by 1D-LCM) in estimating frequency-and-phase drifts and consequentially metabolite concentrations. To this end, we performed Monte Carlo simulations of synthetic in-vivo-like 64-transient MRS experiments with 100 noise realizations at 5 SNR levels with added frequency-and-phase variations and evaluated FPC accuracy, precision, and impact on amplitude estimations. We further tested the proposed method using publicly available in-vivo data from the Big GABA dataset^23^.

## 2. Methods

### 2.1. Synthetic in-vivo-like data

We synthesized realistic in-vivo-like human brain sLASER spectra (TE = 30 ms, 64 transients per dataset) using the MRS data generator that accompanies our open-source analysis toolbox Osprey (https://github.com/schorschinho/osprey)^24^. The generator applies a signal model to metabolite basis functions generated from a fully-localized 2D density-matrix simulation of a 101 × 101 spatial grid (field of view 50% larger than the voxel), implemented in the cloud-based MATLAB toolbox ‘MRSCloud’^25^ which uses FID-A^26^ functions.

We assembled 100 in-vivo-like datasets per SNR level (256, 128, 64, 32, 16), defined as the NAA peak amplitude over one standard deviation (SD) of the noise^27^ in the <u>averaged</u> spectrum, i.e., single-transient SNR was 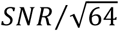. We included 18 metabolite basis functions (ascorbate - Asc, aspartate - Asp, creatine - Cr, CrCH_2_ – a negative creatine methylene accounting for water suppression/relaxation effects, γ-aminobutyric acid - GABA, glycerophosphorylcholine - GPC, Glutathione - GSH, glutamine - Gln, glutamate - Glu, myo-Inositol - mI, lactate - Lac, N-acetylaspartate - NAA, N- acetylaspartylglutamate - NAAG, phosphorylcholine - PCh, phosphocreatine - PCr, phosphorylethanolamine - PE, scyllo-inositol - sI, taurine - Tau) and an experimentally derived macromolecule signal^28,29^ with constant amplitudes and Gaussian (9.7 Hz) and Lorentzian (2.42 Hz) linebroadening.

Each transient received frequency and zero-order phase shifts drawn randomly from normal distributions in the range [-10, 10 Hz] and [-45°, 45°].

## 2.2 In-vivo data

We also tested our new method on publicly available in-vivo human brain data from the Big GABA dataset^23^ (https://www.nitrc.org/projects/biggaba/). Briefly, this dataset included ^1^H MRS PRESS (TE 35 ms; TR 2000 ms; transients 64; spectral bandwidth; 2 kHz or 4 kHz; data points 2074 or 4096; scan time 2 minutes 8 seconds; 30 × 30 × 30 mm^3^ voxel located in the medial parietal lobe) from 73 subjects, acquired at 8 sites using 3 T Siemens scanners (Siemens Healthineers, Erlangen, Germany). The Siemens data were chosen because their .dat/TWIX format preserves the individual transients. We performed coil combination and eddy-current correction on the raw data using the water reference scans followed by HSVD water removal on individual transients and global zero-order phasing based on the Cr and Cho singlets^30^.

The original dataset (73 subjects) had a mean SNR = 269 ± 76, calculated as NAA peak height over one noise standard deviation from the averaged real part of the frequency-domain spectra. We then added random Gaussian noise to each time-domain transient to create five SNR scenarios (again calculated on the averaged spectrum). The amplitude of the added noise was adjusted to approximately correspond to the five SNR levels of the synthetic in-vivo-like dataset. The following mean SNR levels were achieved: 269 ± 76 (original dataset), 135 ± 39, 68 ± 19, 34 ± 10 and 17 ± 5.

### 2.3. Modeling

We modeled all data (synthetic and in-vivo) with a new linear-combination modeling module ^22^ in our MATLAB-based toolbox Osprey^24^, introducing 2D capability (https://github.com/schorschinho/osprey/tree/MSM).

We analyzed each dataset (all SNR levels of synthetic and in-vivo data) with the following two approaches (**Figure 1**):

**Figure 1.**
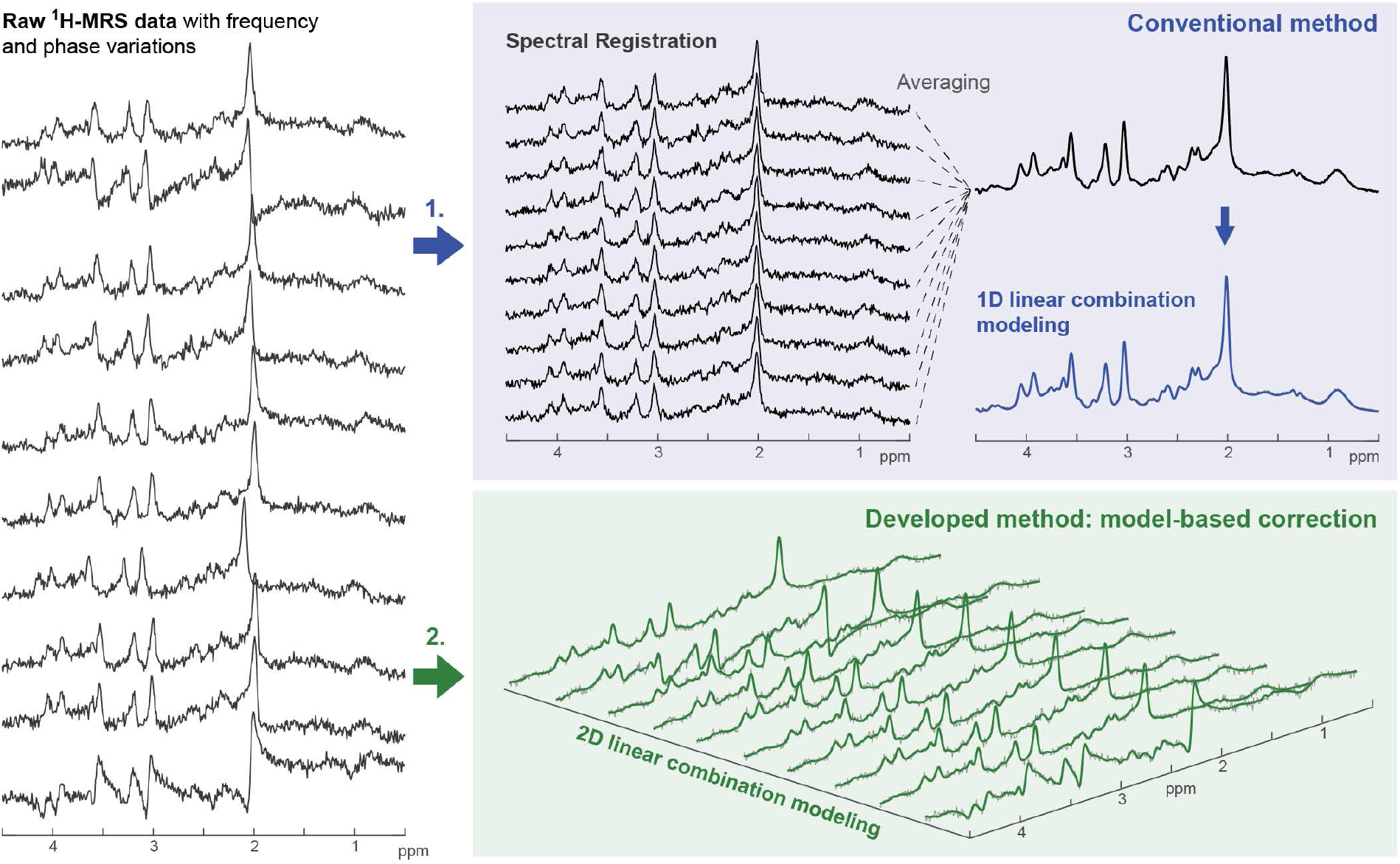
Visualization of the two approaches used for FPC and linear-combination modeling with synthetic realistic in-vivo-like brain spectra. Left panel: Raw MRS data with frequency-and-phase variations. Blue frame (top right): Standard approach with spectral registration followed by averaging and 1D-LCM. Green frame (bottom right): New approach with 2D-LCM performed directly on uncorrected data with individual frequency-and-phase parameters for each transient.

1. “**SpecReg + 1D-LCM**” (traditional approach) FPC with spectral registration^4^, coherent averaging, and 1D-LCM of the averaged spectrum. Spectral registration was performed using the ‘op_alignAverages’ function from FID-A^26^ with the default setting, i.e., all transients are aligned to the single transient with the lowest unlikeness metric, i.e. the one most similar to all other transients. By default, only the first *n* points of the time-domain signal were used for alignment, with *n* the index of the last point before the single-transient time-domain SNR falls below 5. This default cutoff calculation was used for all spectra at the highest SNR levels (on average, the first 115 ms of the FIDs were used for alignment). For lower SNR levels, this criterion disqualified an increasing number of data points used for alignment, until (at SNR = 64 and 32) no points were left. We therefore used the cutoff point determined in the highest SNR level for the lower SNRs (of the same spectrum). As a result, the same number of points in the time domain was used for alignment at every SNR.
2. “**2D-LCM**” Simultaneous 2D modeling of all transients (no prior FPC or averaging) with individual free frequency shift and zero-order phase parameters for each transient. All other model parameters (metabolite amplitude, Gaussian and Lorentzian linebroadening, cubic baseline spline amplitude parameters) were fixed along the transient dimension^22^.

For both approaches, the modeling process was carried out in two steps. In the first step, the metabolite-specific (*local*) frequency shifts were fixed at zero, and a *global* frequency shift and zero-order phase were modeled as free parameters (one each per transient). In the second step, the global frequency and zero-order phase parameters were taken as prior knowledge, i.e., fixed to the values estimated in step 1, with the local frequency shifts now free for each metabolite (but fixed along the transient dimension). This allowed us to avoid competition between global and local frequency shifts. In order to stabilize the solution, our algorithm (similar to the classic LCModel^31^) regularizes certain model parameters by imposing penalty terms of the form 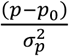, with *p* the estimated value of a parameter, *p*_*0*_ its expectation value, and *σ*_*p*_the SD of its expected distribution. Expectation values (SDs) were 0 Hz (3 Hz) for metabolite-specific frequency shifts 2.75 Hz (2.75 Hz) for Lorentzian linebroadening and 5.12 Hz (5.12 Hz) for global Gaussian linebroadening.

For the synthetic data, least-squares minimization of the difference between the data and model was performed on the real part of the frequency-domain spectrum over a fit range from 0.5 to 4 ppm. Synthetic data were modeled without a baseline component, as no baseline had been added during data generation. The basis sets (18 metabolites and measured MM profiles^28,29^) were the same that were used for data generation^28,29^.

For the in-vivo data, least-squares minimization of the difference between data and model was also performed over a fit range from 0.5 to 4 ppm in the first step. However, in the second step, we introduced a modeling gap between 1.1 and 1.85 ppm to reduce the impact of between-transient fluctuations of difficult-to-model signals (likely brought by phase cycling and varying lipid signals) in that region (**Supplementary Figure 1**). The baseline was parameterized with penalized cubic splines^32^ (P-splines) with a knot spacing of 0.0667 ppm and a fixed regularization parameter Lambda = 15. This regularization approach encourages baseline smoothness and avoids overfitting by the baseline, and is used in other LCM algorithms like ABfit and ProFit^33,34^, similar to the baseline regularization in the original LCModel algorithm^31^. Basis sets included 17 metabolites (lactate was removed from the basis set since its primary resonance at 1.31 ppm was inside the introduced fitting gap) generated as described in section 2.1, with real PRESS pulse waveforms and timings, and corresponding measured macromolecules^28,29^.

### 2.4. FPC evaluation, visualization, and statistics

The performance of the two approaches was evaluated and compared as follows:

#### FPC accuracy and precision

Within each synthetic dataset (64 transients), we plotted the linear regression between the estimated frequency/phase variations and their ground truth and calculated the corresponding R^2^ (used here as a measure of accuracy). The raincloud plots show the distribution of R^2^ at each SNR value for the two approaches where the boxplots included median, interquartile range (IQR), and whiskers (25th or 75th percentile ± 1.5 x IQR). We also calculated the frequency estimation error and phase estimation error, defined as the absolute difference between the true frequency or phase shift and the estimated frequency or phase shift across the 64 transients for each dataset (another measure of accuracy). This will be referred to as absolute error against the ground truth (GTE) in the further text. In addition, we also calculated the SD of the GTE across the 64 transients for each dataset (a measure of precision). This was visualized using boxplots (including median, IQR, and whiskers).

#### Metabolite estimate accuracy, precision, and uncertainty estimation

We evaluated metabolite amplitude estimation by assessing the GTE (to determine accuracy), variances across the number of datasets per SNR level (to determine precision), and relative Cramér-Rao Lower Bounds (CRLB, to determine modeling uncertainty). For synthetic data, we assessed all three quantities; for in-vivo data, which lack a ground truth, we could only determine precision and modeling uncertainty. All metrics were visualized using boxplots (included median, IQR, and whiskers).

#### Statistical testing

We conducted all statistical analyses using R 4.3.1^35^ within RStudio^36^. Mean values of the abovementioned outcome measures were compared between modeling approaches using paired t-tests (R function t.test(X, Y, paired=TRUE)). For cases that did not meet the paired t-test assumptions (normality of difference scores, shapiro.test *p*<0.05), we also conducted a non-parametric paired Wilcoxon Signed-Rank test (<u>R function wilcox.test(X, Y, paired=TRUE).</u> Secondly, non-parametric Fligner-Killeen tests were used to compare variances of metabolite amplitudes and CRLBs between modeling approaches (R function fligner.test(X, Y)).

## 3. Results

All 865 datasets (5 SNR levels × (100 synthetic + 73 in-vivo)) were successfully processed and modeled with the new 2D-LCM and the traditional SpecReg + 1D-LCM approach. Exemplary 1D- and 2D-LCM in-vivo data fits are shown **Figure 2**.

**Figure 2.**
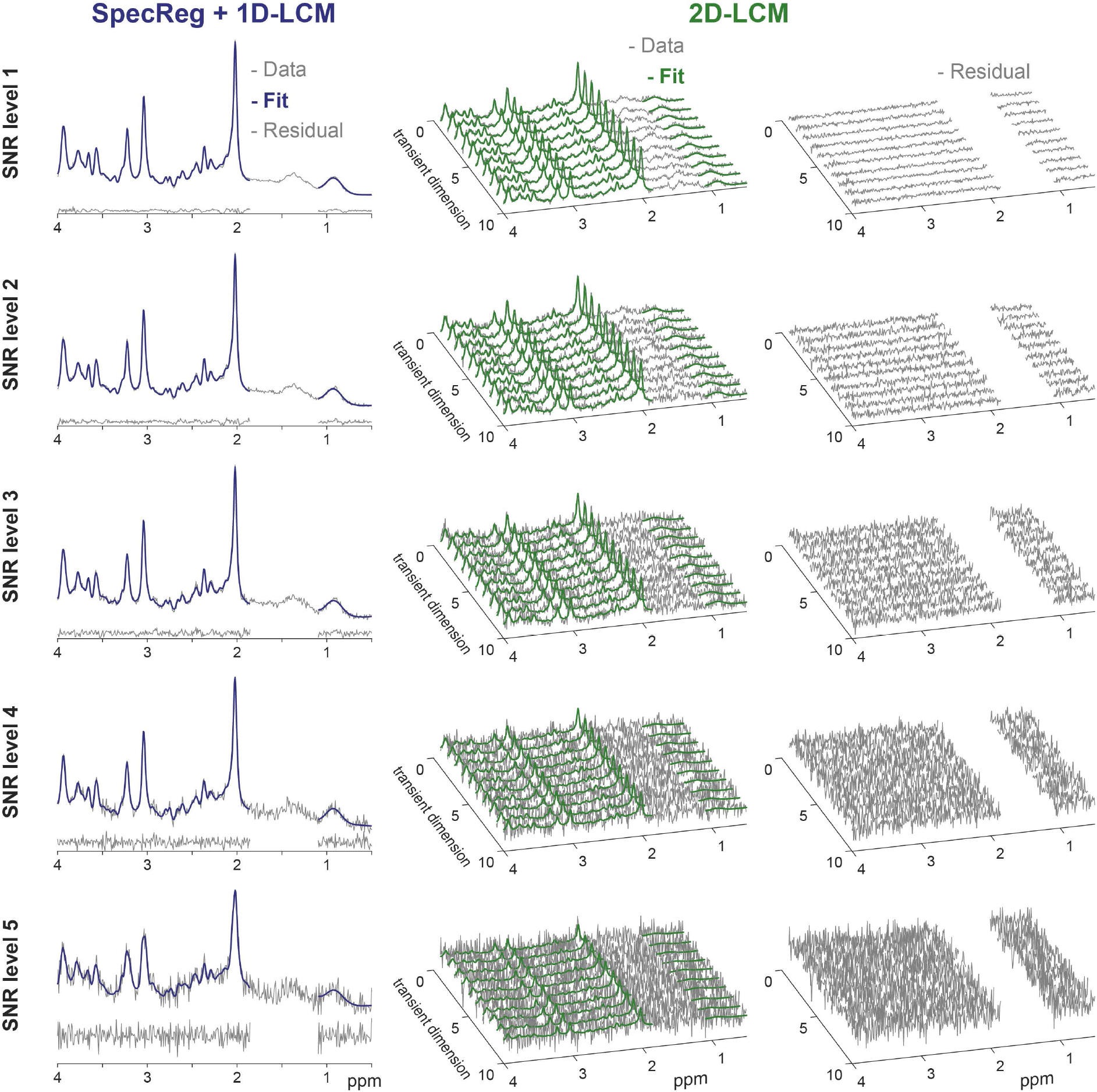
Exemplary 1D- and 2D-LCM in-vivo data fits and the corresponding residuals at five SNR levels. For 1D-LCM, the averaged spectrum is shown with the corresponding fit (in blue) and residual (below). For 2D-LCM a subset of 10 (out of 64) transients are shown with their corresponding fits (in green) and residuals (right panels).

### 3.1. Synthetic data

For synthetic in-vivo-like data at high SNR (256 and 128), SpecReg + 1D-LCM and 2D-LCM both retrieved the frequency and phase with very high accuracy (R^2^ ≥ 0.99) for virtually all datasets (**Figure 3**). The high accuracy was further confirmed with a consistently small GTE (< 1 Hz and < 4.5°) in phase and frequency estimation for SpecReg + 1D-LCM and 2D-LCM. Likewise, both methods achieved excellent precision at high SNR, with the SD of the absolute frequency estimation error consistently below 1 Hz and the SD of phase estimation error below 4.5° (i.e., less than 10% of the maximum shifts), as shown in **Figure 4**.

**Figure 3.**
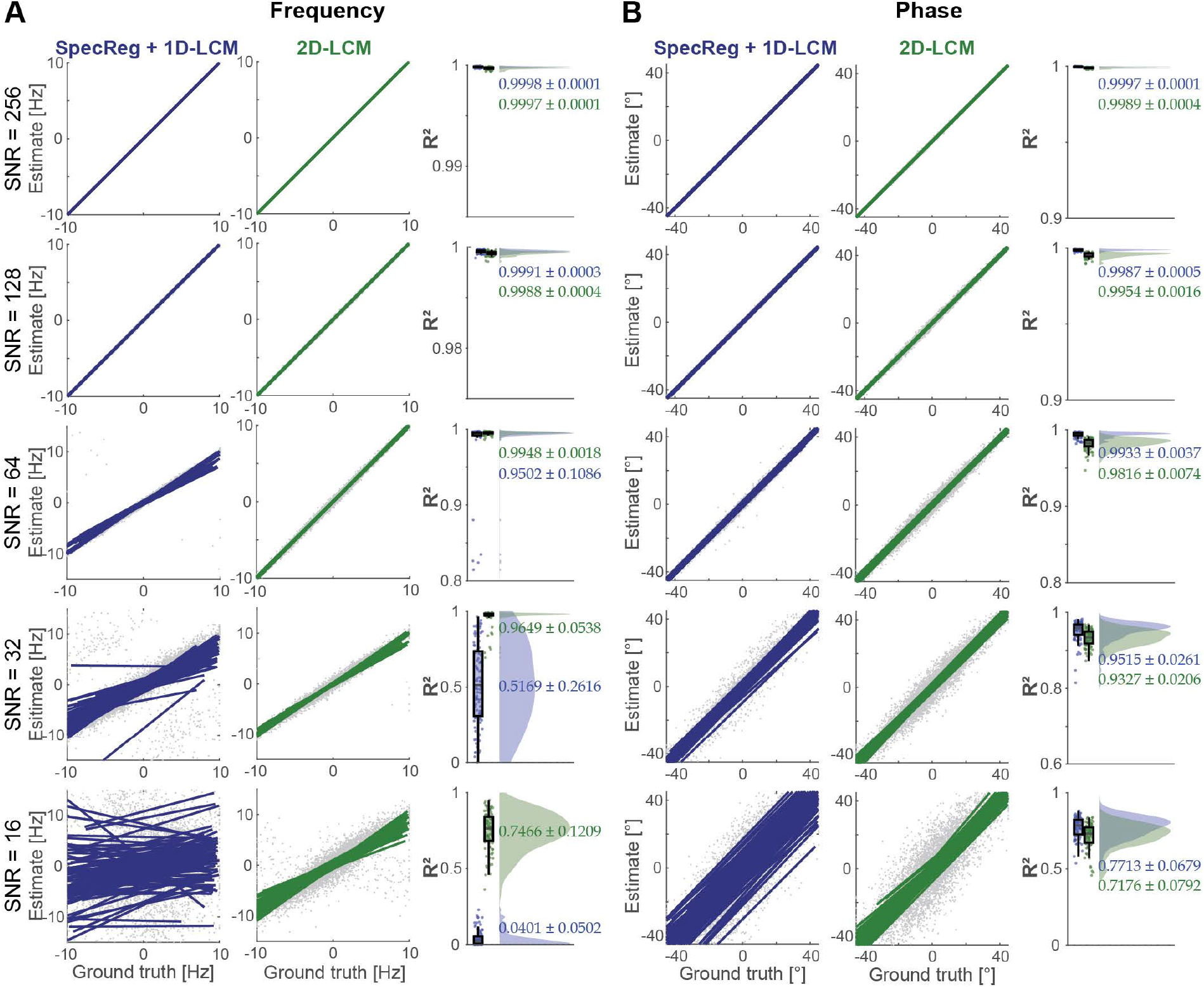
Accuracy of frequency-and-phase shift estimation for SpecReg+1D-LCM approach (blue) and 2D-LCM approach (green) at five SNR levels. The spaghetti plots show 100 linear regressions (on 64 points each for 64 transients per dataset) comparing ground truth and estimates of frequency (A) and phase (B). The corresponding distributions of R^2^ (goodness of fit) are shown in the raincloud plots (right panels) as a measure of accuracy

**Figure 4.**
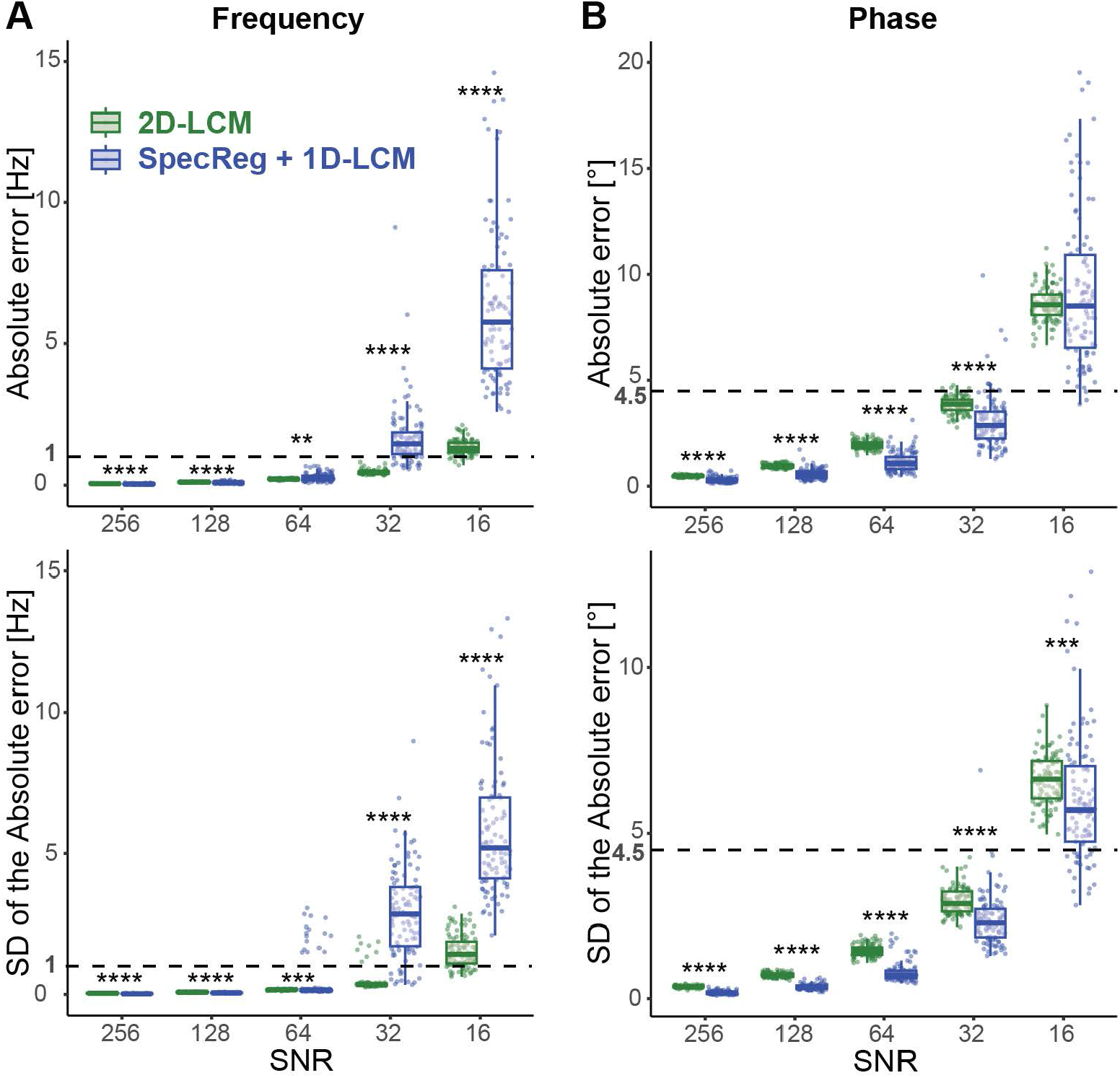
Accuracy and precision of frequency (A) and phase (B) estimation for SpecReg+1D-LCM (blue) and 2D-LCM (green) on synthetic in-vivo-like data. Upper panels show the absolute error against the ground truth (GTE), and lower panels show the SD of the absolute error (SD of the GTE) across the 64 transients per dataset. Box plots represent medians, interquartile ranges, and 25^th^/75^th^ percentile plus/minus interquartile ranges. The dashed black line represents the 10% error mark (1 Hz for frequency and 4.5° for phase). Extreme outliers were excluded from visualization to improve the scaling and clarity of the figure. Means were compared by paired t-test or (in cases where the normality assumption was not met) non-parametric paired Wilcoxon test (**p<0.01, ***p<0.001, ****p<0.0001).

At SNR 64, both approaches maintained excellent accuracy, with an average R^2^ ≥ 0.9 and an average GTE less than 1 Hz and below 4.5°. However, the SD of the frequency GTE increases for SpecReg + 1D-LCM, driven by a group of outliers (**Figure 3&4**), suggesting that single-transient alignment begins to fail entirely for certain datasets at this SNR level, while 2D-LCM maintains excellent precision.

At lower SNR (32 and 16), the difference in performance between the two methods became much more obvious. In general, most metrics for SpecReg + 1D-LCM deteriorated more than for 2D-LCM. In particular, 2D-LCM achieved significantly more accurate frequency shift estimation than SpecReg + 1D-LCM (higher R^2^ and smaller GTE, *p* < 0.0001). Precision was also significantly greater (*p* < 0.0001), as indicated by the SDs of the frequency estimation error.

Both approaches still estimated phase variations very accurately at SNR 32 (R^2^ ≥ 0.9, GTE < 4.5°); while SpecReg + 1D-LCM was (only slightly) more precise. At SNR 16, we observed a decrease in accuracy and precision for both approaches, with GTE and the SD of GTE exceeding the 10% error mark of 4.5° (**Figure 4**). In the case of SpecReg + 1D-LCM, this decrease in performance was less severe than for frequency estimation.

Mean metabolite amplitude estimates agreed well with the ground truth for both approaches at SNR 256 and 128 (**Figure 5**). For both approaches, mean relative CRLBs at SNR 256 and 128 were ≤ 20% for all reported metabolites, indicating that both approaches performed with sufficient certainty at high SNR, although 2D-LCM CRLBs were consistently significantly smaller than 1D-LCM CRLBs.

**Figure 5.**
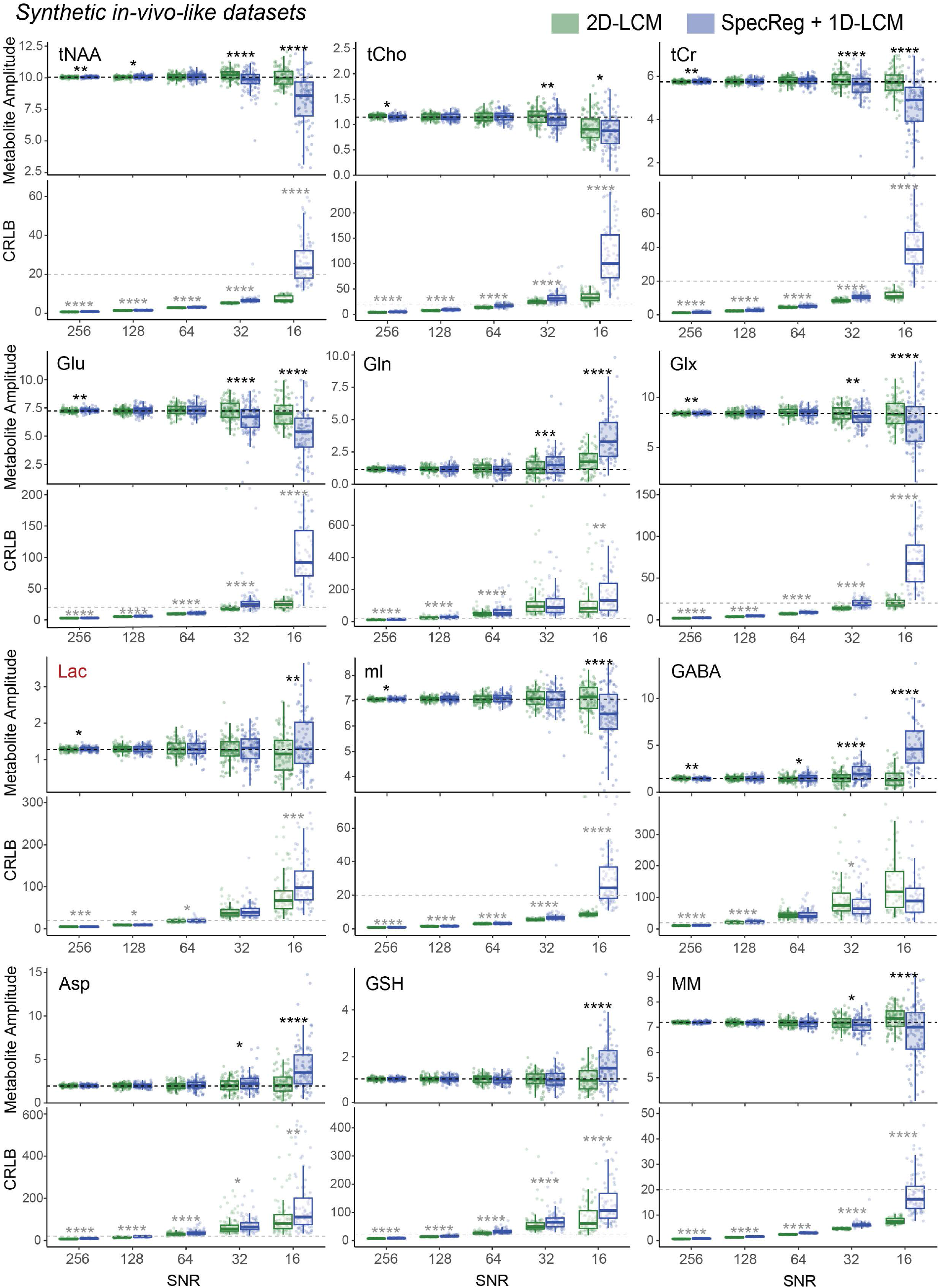
Metabolite amplitudes (upper panels) and relative CRLBs (lower panels) estimated with SpecReg+1D-LCM (blue) and 2D-LCM (green). tCr = Cr+PCr, tNAA = NAA+NAAG, tCho =GPC+PCh; Glx = Glu+Gln. The dashed black line represents ground truth amplitude. The dashed gray line represents 20% CRLB. Infinite CRLBs for zero metabolite estimates were not plotted or considered in statistics. Additionally, some extreme outliers were excluded from visualization to improve the scaling and clarity of the figure. Means were compared by paired t-test or (in cases where the normality assumption was not met) non-parametric paired Wilcoxon test (*p<0.05, **p<0.01, ***p<0.001, ****p<0.0001).

At SNR 64, the mean metabolite amplitude estimates still agreed well with the ground truth. However, relative CRLBs for most of the lower-amplitude metabolites (Gln, Asp, GSH, GABA) exceeded the 20% cutoff point for SpecReg + 1D-LCM and 2D-LCM, although they remained consistently lower for 2D-LCM.

At SNR 32, the high-amplitude metabolites (tNAA, tCr, Glx, and mI) were still accurately estimated by both approaches, although the SD of the tNAA and mI estimates increased substantially for 1D-LCM. Most importantly, relative CRLBs for 2D-LCM remained within 25% for 2D-LCM, but not for 1D-LCM, for Glu (18 ± 3 vs 27 ± 17, *p* < 0.0001) and tCho (25 ± 3 vs 34 ± 13, *p* < 0.0001).

At the lowest SNR level (16), these advantages of 2D-LCM became extremely clear. Amplitude estimates were more accurate for tCr (5.73 ± 0.59 vs 4.67 ± 1.20), Glx (8.43 ± 1.37 vs 7.48 ± 2.33), tNAA (10.02 ± 0.72 vs 8.22 ± 2.09), and mI (7.09 ± 0.52 vs 6.46± 1.08) were closer to the ground truths (5.76, 8.38., 10.05 and 7.06 respectively). They were also more precise (lower SD across datasets) for all metabolites. Finally, 2D-LCM achieved mean CRLBs of ≤ 20% for tCr, tNAA, Glx, and mI even at this lowest SNR level, for which SpecReg + 1D-LCM CRLBs were 2–3 times greater.

In summary, both methods achieved comparable accuracy and precision at very-high-to-moderate SNR. At low-to-very-low SNR, however, 2D-LCM achieved greater accuracy, precision, and modeling uncertainty than the traditional SpecReg + 1D-LCM for all metabolites. SDs of metabolite amplitudes and CRLB estimates are summarized in **Supplementary Figure 2**.

### 3.2. In-vivo data

The overall quality of the original in-vivo data was excellent, with very little between-transient frequency-and-phase variations. The largest range of frequency-and-phase shifts (maximum minus minimum) within a single dataset was 4.5 Hz and 37°, compared to 20 Hz and 90° for the synthetic data. On average, the in-vivo data range of frequency shifts was 1.7 Hz and the range of phase shifts was 15.3°.

For most metabolites, mean amplitude estimates showed only small differences between SNR levels for 2D-LCM. In contrast, they differed substantially for 1D-LCM, especially for the lowest SNR. (**Figure 6** and **Supplementary Figure 3**). This further supports the notion that 2D-LCM maintains in-vivo modeling accuracy even under strongly deteriorating SNR conditions.

**Figure 6.**
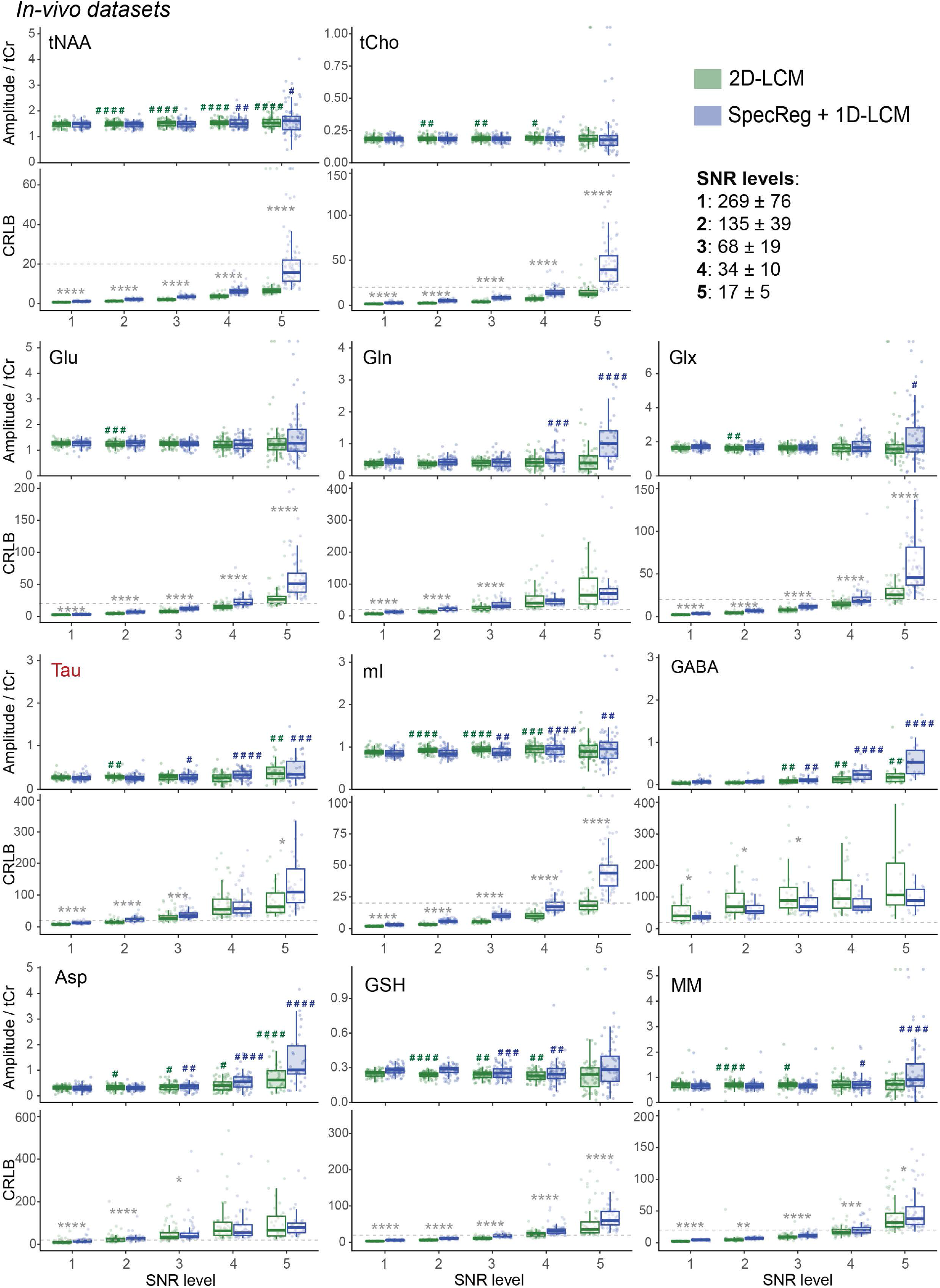
Metabolite amplitudes ratio to tCr (upper panels) and relative CRLBs (lower panels) estimated with SpecReg+1D-LCM (blue) and 2D-LCM (green). tCr = Cr+PCr, tNAA = NAA+NAAG, tCho =GPC+PCh; Glx = Glu+Gln. The dashed gray line represents 20% CRLB. Infinite CRLBs for zero metabolite estimates were not plotted and considered in statistics. Additionally, some extreme outliers were excluded from visualization to improve the scaling and clarity of the figure. Means of the amplitude estimates were compared within each modeling approach, but between SNR levels (always comparing to the highest SNR level), with paired t-tests or non-parametric Wilcoxon test (^#^p<0.05, ^##^p<0.01, ^###^p<0.001, ^####^p<0.0001). Means of the relative CRLB were compared between modeling approaches at each SNR level with paired t-tests or (in cases where the normality assumption was not met) non-parametric paired Wilcoxon test (*p<0.05, **p<0.01, ***p<0.001,****p<0.0001).

SDs of amplitude estimates were comparable between the high SNR levels for both approaches. As SNR decreased, 2D-LCM estimates had significantly lower SDs than 1D-LCM for most metabolites, confirming the improved precision that was also observed in the synthetic data. Again, this effect became very strong for the lowest SNR tier (**Supplementary Figure 4**).

Modeling uncertainty assessment further confirmed the observations in synthetic data. Relative CRLBs were ≤ 20% for all the reported metabolites (except GABA) for both methods at high SNR, although 2D-LCM already achieved slightly, but significantly, lower CRLBs in this SNR regime. At moderate SNR (level 3), CRLBs for most low-amplitude metabolites (Gln, Asp, GSH, Tau) again exceeded the 20% cutoff point for both approaches, but again, they were always lower for 2D-LCM. Finally, at low SNR (levels 4-5), mean CRLBs for tNAA, tCho, Glx, mI, and MM largely remained ≤ 20%, whereas they were much higher (2-3-fold) for SpecReg + 1D-LCM, again consistent with the synthetic data findings (**Figure 6**).

## 4. Discussion

In this work, we present a new method that directly integrates frequency-and-phase correction (FPC) into a 2D linear-combination model (2D-LCM) of all individual transients (‘model-based FPC’). 2D-LCM tools are becoming increasingly popular for dynamic multi-spectrum methods (functional^20^, diffusion-weighted MRS^19^, and relaxometry^37,38^), but as we demonstrate here, they may also offer valuable benefits for conventional (non-dynamic) data^22^. Using synthetic in-vivo-like and in-vivo short-TE datasets, we found that 2D-LCM can estimate (and account for) frequency-and-phase variations directly from uncorrected data with equivalent or better fidelity than the conventional approach, i.e., spectral registration followed by coherent averaging and 1D-LCM. Furthermore, 2D-LCM metabolite amplitudes were at least as accurate, precise, and certain as the conventionally derived estimates, and the advantages of 2D-LCM became strikingly apparent at low to very low SNR. This work suggests that 2D-LCM may hold extraordinary value for data with inherently poor SNR, and may benefit experimental conditions with long TEs, high diffusion weighting^19^, or small voxels.

Alignment of individual transients has become an indispensable processing step for improving spectral quality, particularly for sensitive experiments like spectral editing^39^ and diffusion-weighted MRS^19^. The original time-domain spectral registration algorithm is widely used and incorporated into open-source analysis software^24,26,40,41^. Spectral registration performs fast and efficient FPC but is susceptible to other shot-to-shot signal variations like fluctuating residual water signal, baseline instabilities, or lipid contamination^4,5,16^. More crucially, shot-to-shot alignment becomes challenging in low-SNR conditions^17,42^, as the least-squares optimization requires a certain single-shot SNR level to reliably find the ideal frequency-and-phase shift for an individual transient. Several modifications attempt to address the weaknesses of spectral registration. The RATS algorithm improves robustness to fluctuating baseline signals by incorporating polynomial baseline modeling into variable-projection (VARPRO) optimization in the frequency domain^5^. RATS is generally more accurate and stable for large frequency shifts and unstable baselines, but it is still outperformed by spectral registration at low SNR. Similarly, robust spectral registration (rSR)^16^ attempts to automatically identify transients with lipid contamination or fluctuating residual water, model these signals with a Whittaker smoothing filter, and subtract them out before determining the FPCs. Furthermore, rSR iteratively updates the alignment reference as a weighted average of the already-aligned transients. By maximizing the use of available information from all transients, the updated reference offers better SNR and linewidth compared to the single-transient reference used by the original spectral registration. Both RATS and rSR model parts of the observed signals and attempt to integrate information from all transients into the alignment process, rather than relying on isolated single-transient data. More recently, deep learning FPC algorithms^43,44^ have been introduced, which inherently use all available data as input. In contrast, our 2D-LCM work offers an entirely new perspective on FPC, where the frequency-and-phase shifts in the data are not to be actively corrected for, but rather directly incorporated into a holistic modeling process. 2D-LCM intrinsically maximizes the use of available information across all transients. By assuming that most model parameters like metabolite amplitudes stay constant across the acquisition, the FPC parameters can be estimated with greater accuracy and precision. This gives 2D-LCM surprising stability even in the cases where traditional, non-model-based, spectral alignment fails.

While the conventional spectral alignment methods rely on similarities between spectra for optimal performance, model-based FPC should be feasible even *across* experimental conditions that would prohibit shot-to-shot or spectrum-to-spectrum alignment, e.g., multiple diffusion weightings^4^, edit-ON/OFF^39^. Here, the relationships between spectra can be directly incorporated into the modeling process (e.g., diffusion coefficients, different ON/OFF basis sets, etc.).

Future improvements to model-based FPC should address the influence of shot-to-shot or spectrum-to-spectrum differences that are *not* straightforward to explicitly include in the model. In this work, we encountered inconsistencies between transients in the macromolecular/lipid region between 1.1 and 1.85 ppm, potentially caused by effects of phase cycling on lipids or other unwanted coherence pathways – the source of these fluctuations warrants further investigation. As we were not able to develop a meaningful model for these shot-to-shot fluctuations, the 2D-LCM fits left residuals in the individual transients, while the fluctuations appeared to average out in the 1D-LCM. In this work, we circumvented the problem by introducing a modeling gap that excluded the spectral range of fluctuation. This improved model stability and, as demonstrated above, led 2D-LCM to achieve better modeling performance than 1D-LCM, at the expense of the ability to quantify lactate, and a potential increase in the uncertainty of GABA estimation, as the 1.89 ppm ^3^CH_2_ signal is close to the gap. For true universal applicability to in-vivo data, 2D modeling may need to consider the inconsistencies between transients. These artifacts may include out-of-voxel echoes, lipid signals, different phase cycling steps, etc. Alternatively, 2D-LCM may still offer substantial benefits if only applied to averaged phase cycles instead of individual transients, which may remove most of the fluctuations. Finally, gross outlier transients (e.g., motion corruption) may require pre-identification and downweighing to not skew the overall 2D model.

2D modeling can be biased by the incorrect selection of spectral models that do not describe data behavior correctly^21^ — either in the spectral dimension or the second, indirect dimension^45^. It is well-known that the choice of spectral model, modeling parameters, starting values, upper/lower bounds, regularizations, and penalties has an outsized impact on modeling results, even for 1D-LCM^30,46–51^. The impact of parameters that characterize the model along the second dimension is less well investigated but should be the focus of future studies to ascertain that 2D-LCM is indeed more accurate and precise (and not just more rigid). Regularization strategies are frequently used in linear-combination modeling to constrain the solution space of the ill-defined optimization problem to meaningful values but are currently mostly applied to metabolite or baseline amplitude parameters^31,33^. They may also be very useful in 2D-LCM and could be tailored for specific purposes. In the example of FPC, thermal frequency drift usually follows a smooth trajectory in time. Applying another regularization parameter to impose smoothness on the frequency and phase shift parameters in their order of acquisition may then further stabilize (and accelerate) 2D-LCM and model-based FPC.

Finally, future work should also explore whether model-based FPC can be directly incorporated into more dynamic and complex 2D modeling scenarios like diffusion-weighted, functional, edited, hyperpolarized MRS, and relaxometry^45^. Recent studies of 2D modeling^21,45^ have generally found improvements in fitting uncertainty and precision of target parameter estimation in several dynamic data scenarios. Certain benefits of 2D-LCM in uncertainty estimation were also shown for conventional 1D-MRS data^22^. With 2D-LCM becoming increasingly available in open-source analysis software, their respective algorithms should be able to evolve and adopt new functionality in the coming years^24,40,52–58^.

## Conclusion

Model-based FPC with 2D linear-combination modeling is feasible and outperforms traditional transient-to-transient alignment methods for lower SNRs.

## Acknowledgments

This work has been supported by NIH grants R00 AG062230, R21 EB033516, K99 AG080084, K00AG068440, R01 EB016089, R01 EB023963, R01 EB032788, and P41 EB031771.

