## Supplementary Material for "Model-based frequency-and-phase correction of ^1^H MRS data with 2D linear-combination modeling"

### Synthetic in-vivo-like data

To generate synthetic in-vivo-like data the following amplitudes were given to the basis functions Asc - 0.19, Asp - 1.94, Cr - 3.13, CrCH2 - 2.14, GABA - 1.46, GPC - 0.54, GSH - 1.07, Gln - 1.15, Glu - 7.23, mI - 7.06, Lac - 1.28, NAA - 8.70, NAAG - 1.35, PCh - 0.61, PCr - 2.63, PE - 1.72, sI - 0.13, Tau - 0.39, MM - 7.2.

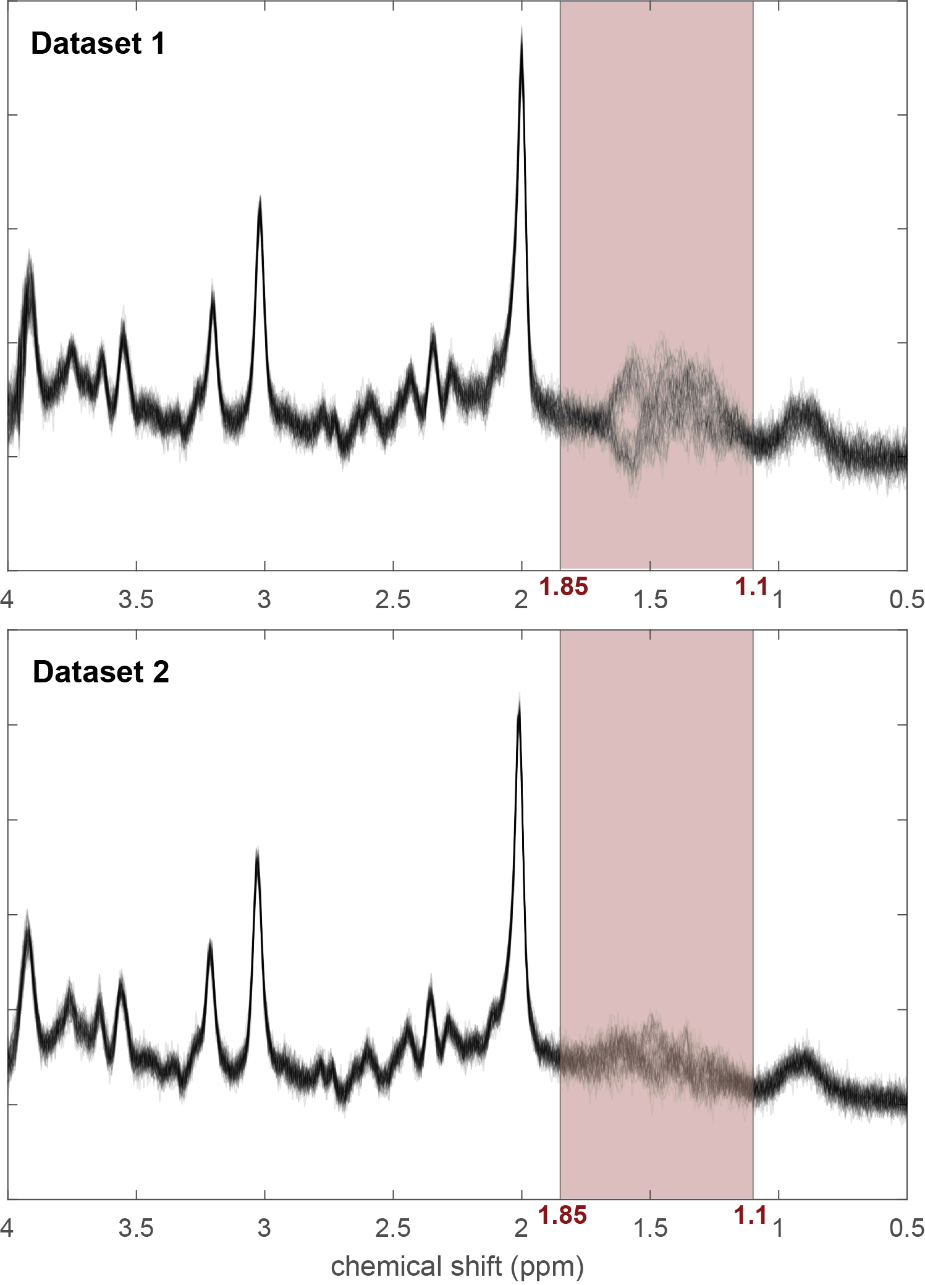

*Supplementary Figure 1. Two example in-vivo datasets (overlay of 64 transients). Red shading highlights the area between 1.85 and 1.1 ppm where the between transient fluctuations (likely brought by phase cycling and varying lipid signals) appear.*

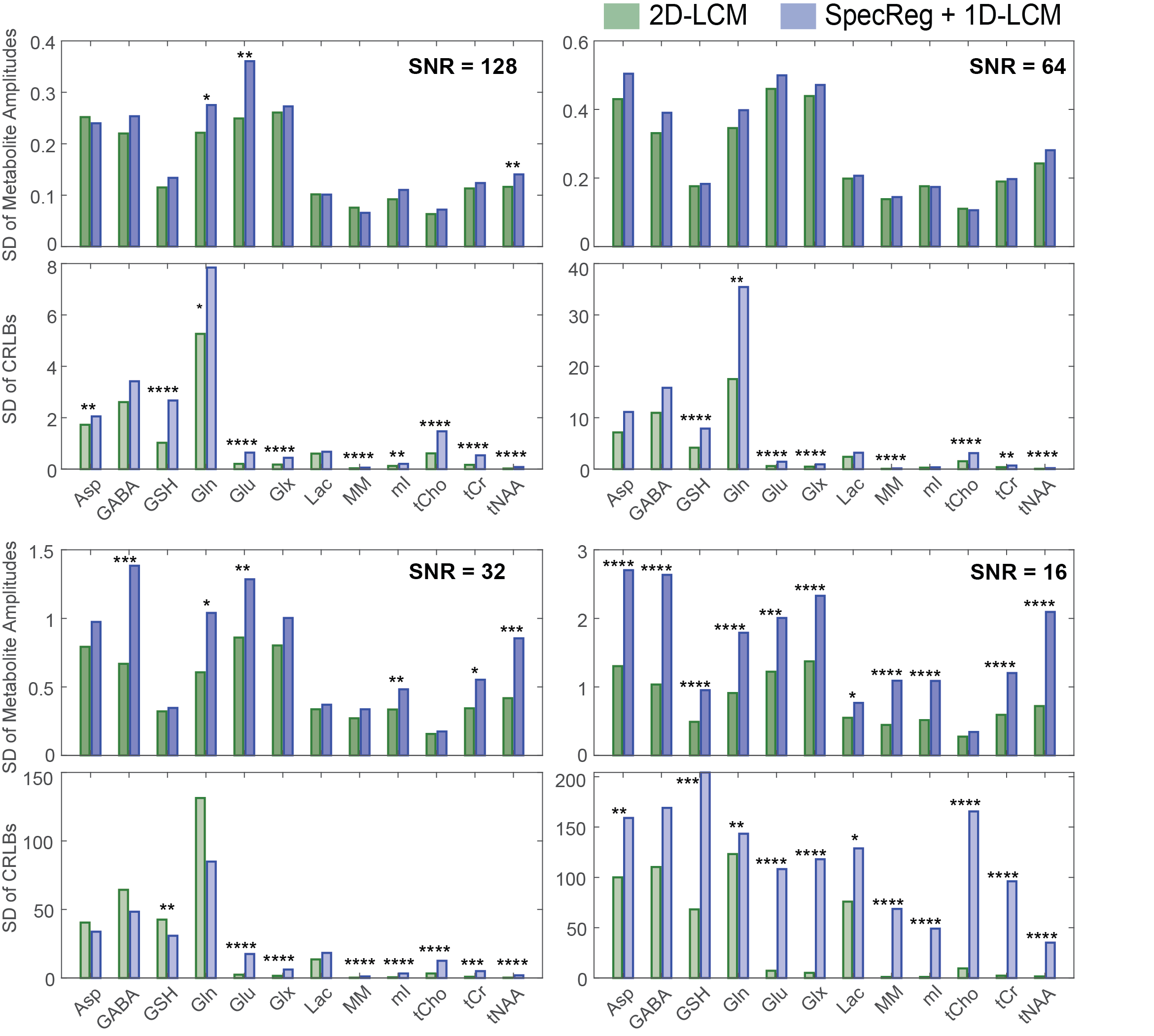

*Supplementary Figure 2. Synthetic in-vivo-like datasets. Standard deviations of metabolite amplitudes (top panel) and the corresponding CRLBs (bottom panel) at SNR 128, 64, 32 and 16. Non-parametric Fligner-Killeen tests were used to compare variances of amplitudes and CRLBs for each metabolite (*p<0.05, **p<0.01, ***p<0.001, ****p<0.0001).*

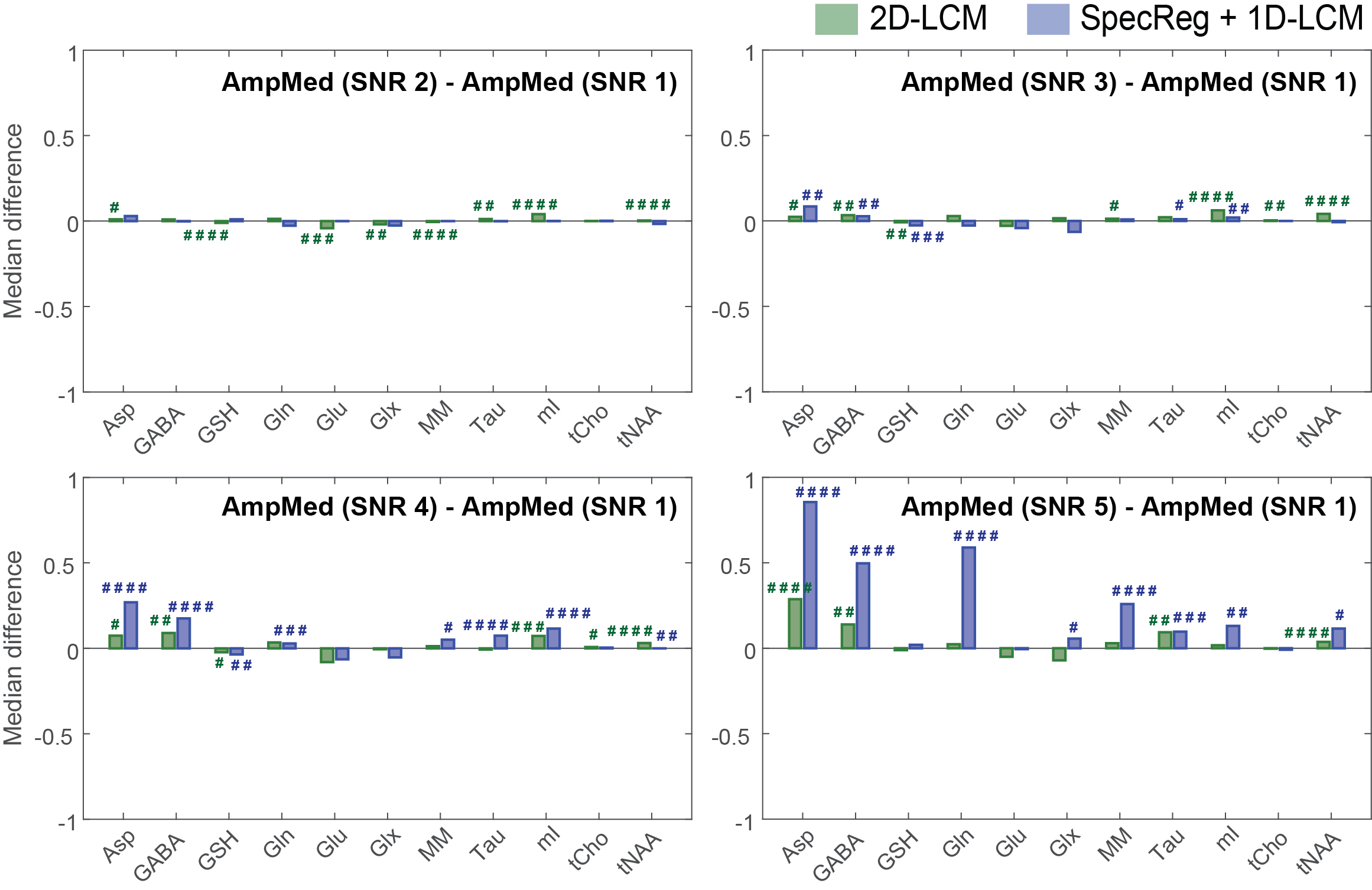

*Supplementary Figure 3. Differences between medians of in-vivo amplitude estimations (AmpMed) at highest SNR level (SNR 1) and all the lower SNRs (SNR levels 2, 3, 4 and 5). The difference is always calculated as AmpMed at SNR X – AmpMed at SNR 1. The statistics show the t-test comparisons of the amplitude estimates within each modeling approach, but between SNR levels (always comparing to the highest SNR level), with paired t-tests or (in cases where the normality assumption was not met) non-parametric paired Wilcoxon test (# = p<0.05, ## = p<0.01, ### = p<0.001, #### = p<0.0001) – same as Figure 5.*

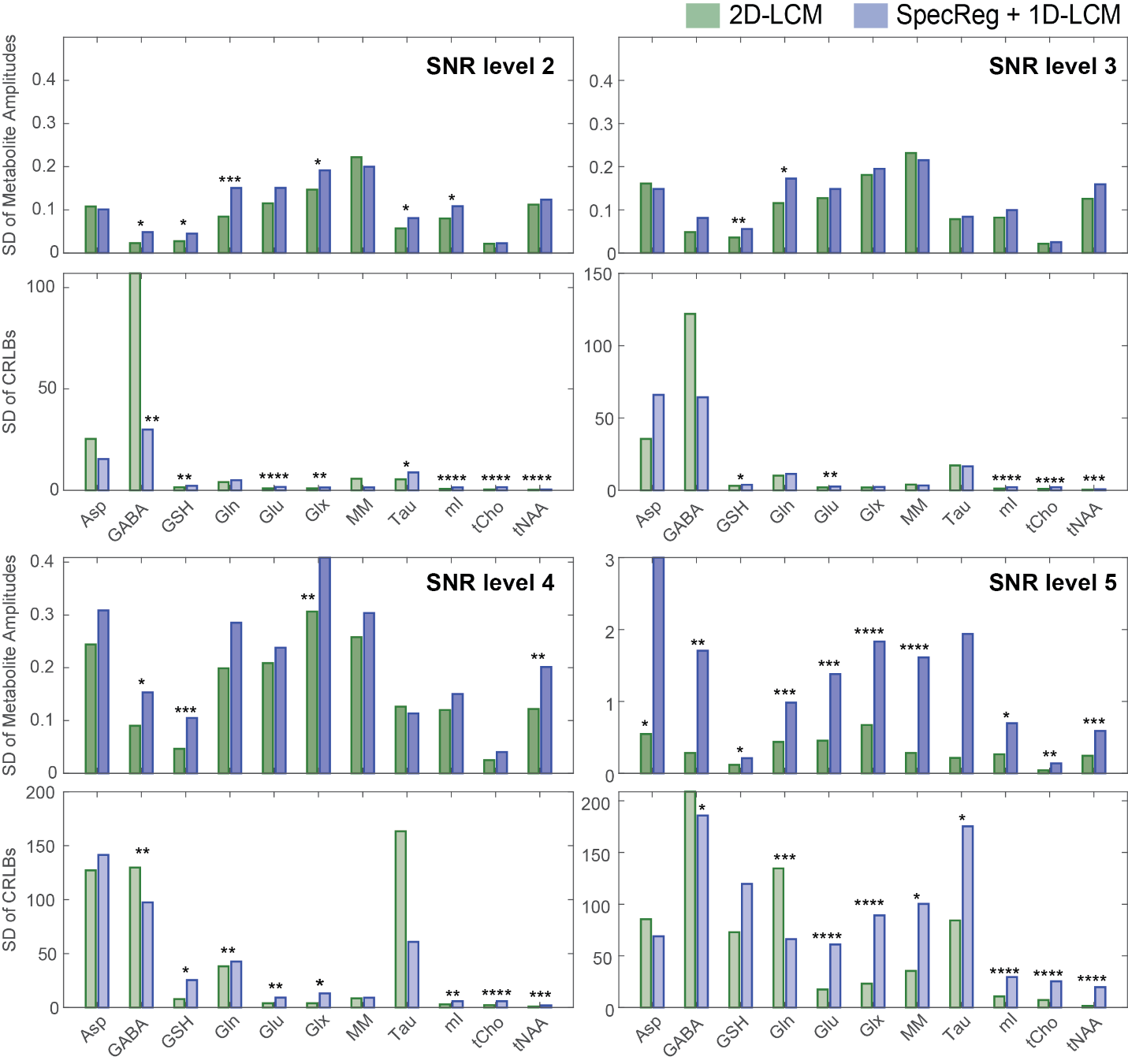
*Supplementary Figure 4. In-vivo datasets. Standard deviations of metabolite amplitudes (top panel) and the corresponding CRLBs (bottom panel) at SNR levels 2, 3, 4, and 5. Non-parametric Fligner-Killeen tests were used to compare variances of amplitudes and CRLBs for each metabolite (*p<0.05, **p<0.01, ***p<0.001, ****p<0.0001).*
